# Brain large artery dilatation increases the risk for Alzheimer’s disease pathology

**DOI:** 10.1101/2025.08.29.672901

**Authors:** Dominic Simpson, Christopher D. Morrone, Darcy Wear, Adrian Khani, Fang Liu, Jose Gutierrez, Wai Haung Yu

**Affiliations:** Department of Pharmacology and Toxicology, University of Toronto, Toronto, Canada; Brain Health Imaging Centre, Centre for Addiction and Mental Health, Toronto, Canada; Department of Neurology, Columbia University, New York, NY, United States

## Abstract

Alzheimer’s disease (AD) and related dementia cases are increasing globally, emphasizing the urgent need to clarify disease mechanisms for translational application in diagnoses and treatment. Vascular alterations represent a major pathological feature of AD, and beyond the well-established roles of small vessel disease and large artery atherosclerosis, our group has previously demonstrated that brain large artery dilatation is associated with elevated risk of dementia and Alzheimer pathology. The most severe manifestation of this non-atherosclerotic arterial phenotype is dolichoectasia, an enlargement of large blood vessels (Gutierrez et al., 2019; Melgarejo et al., 2024). Despite consistent epidemiological evidence across populations, the mechanistic link between arterial dilatation and AD remains poorly understood.

To address this gap, we induced dolichoectasia in *App*^NL-G-F^ mice, a model of amyloid pathology, by injecting elastase into the cisterna magna. After three months, brains were examined using biochemical and immunohistochemical methods. Elastase-treated mice exhibited a significant increase in amyloid plaques in the hippocampus (**p = 0.021**) and cortex (**p = 0.029**) compared with vehicle-treated controls. Neuronal loss was evident in the CA1 region of the hippocampus (**p = 0.036**), with a trend towards neurodegeneration in CA3 (p = 0.055). We also observed elevated p62 in the hippocampus and cortex (**p = 0.009** and **p = 0.001**, respectively), suggesting impaired protein or autophagic-lysosomal clearance. Although no overt increase in neuroinflammation or astrogliosis was detected at this time point, matrix metalloproteinase-9 (MMP-9) levels were trending towards elevated levels (p = 0.058).

Combined, these findings indicate successful elastase-induced brain arterial dilatation accelerates AD-related pathology in *App*^NL-G-F^ mice, providing mechanistic evidence that large artery dilatation may contribute directly to Alzheimer’s disease progression.

## BACKGROUND

Alzheimer’s disease (AD) is the leading cause of dementia worldwide, pathologically defined by extracellular amyloid plaque deposition, intracellular neurofibrillary tangles, chronic neuroinflammation, and progressive neuronal loss in regions critical for cognition (DeTure & Dickson, 2019). Vascular dementia (VaD) represents the second most common dementia subtype and is characterized by cerebrovascular pathology that restricts cerebral blood flow, leading to hypoxia, neuronal death, and cognitive decline (Duong, Patel, & Chang, 2017). Both AD and VaD share modifiable vascular risk factors—including hypercholesterolemia, diabetes, hypertension, and stroke history—and increasing evidence suggests overlapping mechanisms, particularly the interplay between vascular impairment and amyloid deposition (Duong, Patel, & Chang, 2017; Livingston et al., 2024). This interaction underlies the frequent occurrence of mixed dementia and the rising co-morbidity of AD and VaD with advancing age (Barker et al., 2002; Cortes-Canteli & Iadecola, 2020).

The contribution of vascular dysfunction to AD pathogenesis are not fully understood. Clinical and postmortem studies provide important correlative evidence but are limited in their ability to provide mechanistic insight. Consequently, preclinical approaches are essential to provide insight into how vascular alterations influence AD pathology. Of particular interest is brain large artery dilatation—a non-atherosclerotic vascular phenotype characterized by loss of arterial wall elasticity and morphological remodeling—which can impair cerebral blood flow and increase dementia risk in humans (Del Brutto, Ortiz, & Biller, 2017)(Gutierrez et al., 2019).

Experimental models have successfully induced dolichoectasia, the extreme form of arterial dilatation, in rodents, demonstrating its effects on cerebrovascular function and providing an avenue to interrogate its role in neurodegeneration (Dai et al., 2015; Liu et al., 2022; Simpson et al., 2024).

Arterial dilatation can compromise the blood-brain barrier (BBB) through hypoxia-driven astrocytic responses (de la Torre & Mussivand, 1993) and degradation of the vascular elastin lamina, leading to upregulation of vascular endothelial growth factor (VEGF) and consequent pathological angiogenesis and vascular leakiness (Desai, Schneider, Li, Carvey, & Hendey, 2009; Gutierrez, Sacco, & Wright, 2011). Although compensatory upregulation of tight junction proteins may occur, these processes can paradoxically weaken BBB integrity Biron, Dickstein, Gopaul, & Jefferies, 2011). Increased permeability facilitates the entry of peripheral toxins and may exacerbate amyloid plaque deposition, amplifying neuroinflammation, neuronal loss, and cognitive impairment (Zenaro, Piacentino, & Constantin, 2017). For this study, we test the hypothesis that brain large artery dilatation accelerates AD pathology using the *App*^NL-G-F^ mouse model of amyloid deposition, examining the mechanism of neurodegeneration as related to canonical pathologies, proteostasis and neuroinflammation.

## METHODS

Twenty *App*^*NL-G-F*^ mice (12 male and 8 female) at 6 months of age were used in this study. The use of animals and the study protocol were reviewed and approved by the Animal Care Committee (Protocol #843) at the Centre for Addiction and Mental Health, Toronto, Canada. *App*^*NL-G-F*^ mice were outbred in-house and housed on a 12-h light- dark schedule with *ad libitum* access to chow and water.

### Elastase injection

Injection was performed as previously described in our methodological paper (Simpson et al., 2024). Six months (6) old mice were anesthetized with isoflurane (5% induction, 2.5-3% maintenance mix with 2% oxygen), and given local anesthetic (bupivacaine) and analgesic (Metacam). Saline was administered subcutaneously to the animal to keep it hydrated.

The animal’s head was shaved at the base of the skull and positioned in a prone position on a stereotaxic frame while maintaining anesthetic plane via delivery through the specialized nose cone. A bar was added to the top of the mouse’s head to provide stability, and the upper front teeth were fastened. After that, the head was bent at about 120° angle, which elevated and distended the nape. The region was cleaned with betadine scrub (three swipes), 70% ethanol (three swipes), and betadine solution (one swipe) using sterile gauze. For separating the muscles to expose the skull, the midline of the nape of the neck was incised. After separating the muscles with a pair of tweezers to reveal the base of the skull, the cisterna magna (inverted triangle) was located beneath the base of the skull (**Supplemental Figure 1**). The Hamilton syringe’s bevel was slowly inserted into the center of the cisterna magna, avoiding puncturing any blood vessels that ran across the CM. Following the CM puncture, the bevel was removed to facilitate the release of a small amount of cerebrospinal fluid (CSF), and the extra CSF was absorbed using a cotton Q-tip. The bevel was then reinserted at the puncture site, 2.5 µL of elastase (concentration of 15 mU) was slowly injected into the CM, and the needle was left in place for 1 minute (to prevent elastase leaking) before being removed. Sutures were used to close the wound. After being revived from anesthesia, the mice were given a post-surgical injection of Metacam (analgesic)(Simpson et al., 2024).

### Perfusion of Mice

At 9 months of age the animals were administered an overdose of avertin (1ml, 125-250 mg/kg) to induce deep anesthesia. The mouse was perfused using PBS containing 1% heparin to clear blood vessels. Afterward, the brain was extracted, the left hemisphere was dissected for biochemical analysis, while the right hemisphere was post-fixed in 4% PFA at room temperature for immunohistochemistry. The subsequent day, the post-fixed hemisphere was washed in PBS (3 times), followed by immersion in 30% sucrose solution and stored at 4°C.

### Brain Sectioning and Immunofluorescence staining

#### Brain sections

post-fixed brains were sectioned at 40 µm coronally on a microtome (Leica SM2000R) and stored in cryoprotectant at -20°C. Two sections spaced 10 apart were sampled throughout the hippocampus (starting: Bregma -2.155 mm). The sections were placed in a 24-well plate and washed thrice with PBS. For the Iba1 staining, antigen retrieval was performed by placing the sections in sodium citrate solution (pH 6) within a 24-well plate, followed by an 80°C incubation for 20 minutes. Post-incubation, they were rinsed three times with PBS to remove excess sodium citrate. To prevent non-specific binding, a 1 hr incubation at room temperature was done using a blocking solution comprising 2% goat serum [ThermoFisher, Canada, 16210072], 0.1% Triton-X 100 [Bioshop, Canada, 9002-93-1], and 1% BSA [Bioshop, Canada, 9048-46-8], all prepared in PBS.

#### Immunostaining

Primary antibodies used included: rabbit anti-GFAP with a 1:500 dilution [Dako, Glostrup, Denmark; Z0334, polyclonal], rabbit anti-IBA1 with a 1:1000 dilution [Wako, Neuss, Germany; 019-19741, polyclonal], mouse anti-NeuN with a 1:500 dilution [Millipore Sigma, Oakville, Canada; MAB377, monoclonal] and rabbit anti-p62 with a 1:400 dilution [Abcam, Cambridge, United Kingdom; Ab109012, polyclonal]. For the secondary antibodies, goat anti-rabbit 488 with a 1:200 dilution [Invitrogen, Massachusetts, United States; A11008, polyclonal], goat anti-mouse 568 with a 1:200 dilution [ThermoFisher, Massachusetts, United States; A11004, polyclonal], and goat anti-rabbit 568 with a 1:200 dilution [ThermoFisher, Massachusetts, United States; A11075, polyclonal]. For staining of amyloid plaques, Thioflavin-S (ThioS) dye [Millipore Sigma, Oakville, Canada; T1892] was used.

The same blocking solution used to dilute the primary antibodies was added to each incubating chamber and maintained overnight at 4°C shielded from light. The following immunostaining was conducted - GFAP/NeuN, IBA1 and p62. After the primary incubation, the sections were washed three times with PBS, and the respective secondary antibodies (matched to the host of the primary antibody) were added to each well. Incubation was done for 2 hours at room temperature, shielded from light by foil. The secondary antibodies used included anti-rabbit 488/anti-mouse 568 (GFAP/NeuN), anti-rabbit 488 (p62) and anti-rabbit 568 (Iba1).

After the secondary incubation to detect the primary antibodies, the sections were washed three times with PBS and prepared for mounting. An additional step was carried out for the Iba1 stain, where Thioflavin S dye was made up (1% wt/vol ddH2O then syringe filtered) and applied to each section for 7 minutes to label the beta-sheet structure of amyloid plaques. Subsequently, the sections underwent two washes with 70% ethanol <u>(Sigma)</u> for 5 minutes each, followed by three washes with PBS. Sections were mounted and coverslipped in ProLong™ Gold Antifade Mountant (ThermoFisher, P36930). See Supplementary Table 1 for vendor and catalog information.

#### Imaging

Fluorescent images were captured at 10x using the Olympus VS200 slide scanner and VSI software. The captured images were analyzed using the software ImageJ where the images were processed, binarized, and analyzed for staining density (% area covered and number of cells/mm2).

### Biochemical Analysis /Western Blot

#### Sample homogenization

Dissected brain tissues were weighed to calculate the amount of homogenization buffer to be used. A Pierce protease and phosphatase inhibitor (Sigma Aldrich, Canada, P8340), 100mM phenylmethylsulfonyl fluoride (PMSF, Millipore Sigma), 100mM sodium fluoride (NAF, Millipore Sigma), 100mM sodium orthovanadate (Na3V04, Sigma), and 100mM EGTA (Sigma) was used at a 1:100 ratio in Radioimmunoprecipitation Assay buffer (RIPA containing 50 mM Tris-HCl (pH 7.4), 150 mM NaCl, 1% Triton X-100, 0.5% Sodium Deoxycholate (Millipore Sigma) and 0.1% Sodium Dodecyl Sulfate (Millipore Sigma)) as the homogenization buffer. 1mg of tissue was combined with 1mL of chilled homogenization buffer and sonicated at a 30% duty cycle for 5 seconds on wet ice, repeated twice, with the sonicator tip rinsed with distilled water between each cycle. The samples were then centrifuged at 5000 x g for 15 minutes at 4°C. The supernatants were transferred to clean tubes and stored at -80°C until used. A protein assay (Bicinchoninic acid assay) was used to quantify total protein concentration in the supernatant (Pierce™ Protein Assay Kits, ThermoFisher Scientific, Catalog: 23227). For electrophoresis, samples containing 30 μg total protein for MMP-9 detection and 10 μg for PHF1 (Ser396/Thr404 phosphorylated Tau) were prepared by mixing with Dithiothreitol (DTT) (1:10 1M stock solution (ThermoFisher, R0861), LDS 4x dye (ThermoFisher), and distilled water. The mixture was then electrophoresed using 4-20% tris-glycine polyacrylamide gels (ThermoFisher) under reducing conditions.

After electrophoresis, the gel was then transferred using tris-glycine transfer buffer containing 25 mM Tris and 192 mM glycine and 20% methanol, onto 0.2uM nitrocellulose membrane (Bio Rad, 1620112). The transfer was conducted at 200 mA for 2 hours at room temperature with a cold pack to reduce heat. Following transfer, the membranes were immersed in Ponceau S solution for 5 minutes and rinsed with water to confirm consistent protein transfer. Membranes containing Ponceau S staining were digitally captured using the Cytiva Image Quant 800 (Amersham). Following this, the blots were blocked with 5% milk in Tris-buffered saline 0.05% Tween 20 (TBS-T) for 1 hour at room temperature. Rabbit anti-MMP-9 (Proteintech, Rosemount, United States; 10375-2-AP, polyclonal) and Mouse anti-PHF1 (gift from Peter Davies, monoclonal) antibodies were diluted (1:1500 and 1:1000 respectively) in SuperBlock/TBS Solution and left to incubate with the blots overnight at 4°C. For the control, Mouse anti-GAPDH (Millipore Sigma, Oakville Canada; G8795) was diluted at 1:4000 in SuperBlock/TBS Solution. Horseradish peroxidase-conjugated goat anti-mouse IgG for PHF1 and GAPDH (1:10000, 1 hour, Jackson Immunoresearch Labs, West Grove, PA) and horseradish peroxidase-conjugated goat anti-rabbit IgG for MMP-9 (1:10000, 1 hour, Jackson Immunoresearch Labs, West Grove, PA), were used along with the ECL detection system to detect bound antibodies. The blots were imaged using the Cytiva Image Quant machine (Amersham ImageQuant™ 800) and the Densitometric analysis for MMP-9 (67 kDa) and PHF1 (56-65 kDa) specific bands was performed with GAPDH as a control using ImageJ.

### Statistical analysis

All results are shown as group means [± standard deviation (SD)]. An unpaired t-test was used to compare the means between the two independent groups. We also performed a two-way analysis of variance (two-way ANOVA), followed by a post hoc Holm-Sidak test (*α* < 0.05). The GraphPad Prism 10 was used for statistical analysis where p values < 0.05 were considered statistically significant.

## RESULTS

In this study, we successfully induced dolichoectasia in the *App*^*NL-G-F*^ mouse model of AD by injecting elastase in the magna cisterna, to investigate the relationship between vascular impairments and AD pathology. We selected this model due to its ability to form amyloid plaques, a pathology starting early in AD, while expressing the mutant form of human amyloid precursor protein (APP) at physiological levels (Saito et al., 2014). As originally described, amyloid plaques form as early as 4 months, and reach maximal load by approximately 12-16 months, with memory impairment at 6 months (Saito et al., 2014). To induce brain large artery dilatation, we injected elastase into the cisterna magna located above the Circle of Willis.

Elastase, known for its ability to degrade elastin, disrupts the BBB, and is an effective method for inducing brain large artery dilatation in mice (Dai et al., 2015; Takata et al., 2015; Temesvari et al., 1995) and which we recently validated (Simpson et al., 2024).

### Dolichoectasia increased amyloid plaque deposition but did not induce overt neuroinflammation

We investigated the role of brain large artery dilatation in promoting amyloid plaque production by conducting Thioflavin-S staining on the hippocampus the App^*NL-G-F*^ mice after elastase injection (**Fig. 1A and 1B**). Our findings revealed a significantly higher deposition of amyloid plaque in the hippocampus of elastase-injected mice relative to vehicle (PBS-treated) mice [t (18) = 2.540, **p=0.021** and F (1,16) = 8.753, **p=0.009]**; **Fig. 1C**. There was an overall sex difference observed, where both elastase and vehicle female App^*NL-G-F*^ mice which had a higher amyloid deposition than the male counterparts [F (1,16 =6.370, **p=0.023**]. Representative images of the CA3 region of the hippocampus at x10 magnification demonstrate larger plaques in elastase-injected mice (**Fig. 1Ai and 1Bi**). Even though there was an increase in amyloid deposition in the hippocampus, there was no increase in Iba1 activation (**Fig. 1D**). Results in the cortex were similar to the hippocampus (**Fig. 1E**), including a significant increase in plaque deposition after elastase injection (t (18) = 2.378, p=0.029 & F (1,16) = 7.358, **p=0.015** (**Fig. 1F**) and an overall sex difference with higher plaque-loads in both the elastase and vehicle female [F (1, 16) = 17.96, **p= 0.0006**]. However, a significant difference was observed in the male *App*^*NL-G-F*^ mice treated with elastase compared to the vehicle treated group, suggesting that large vessel dilated mice are more susceptible to amyloid plaque deposition.

**Figure 1:**
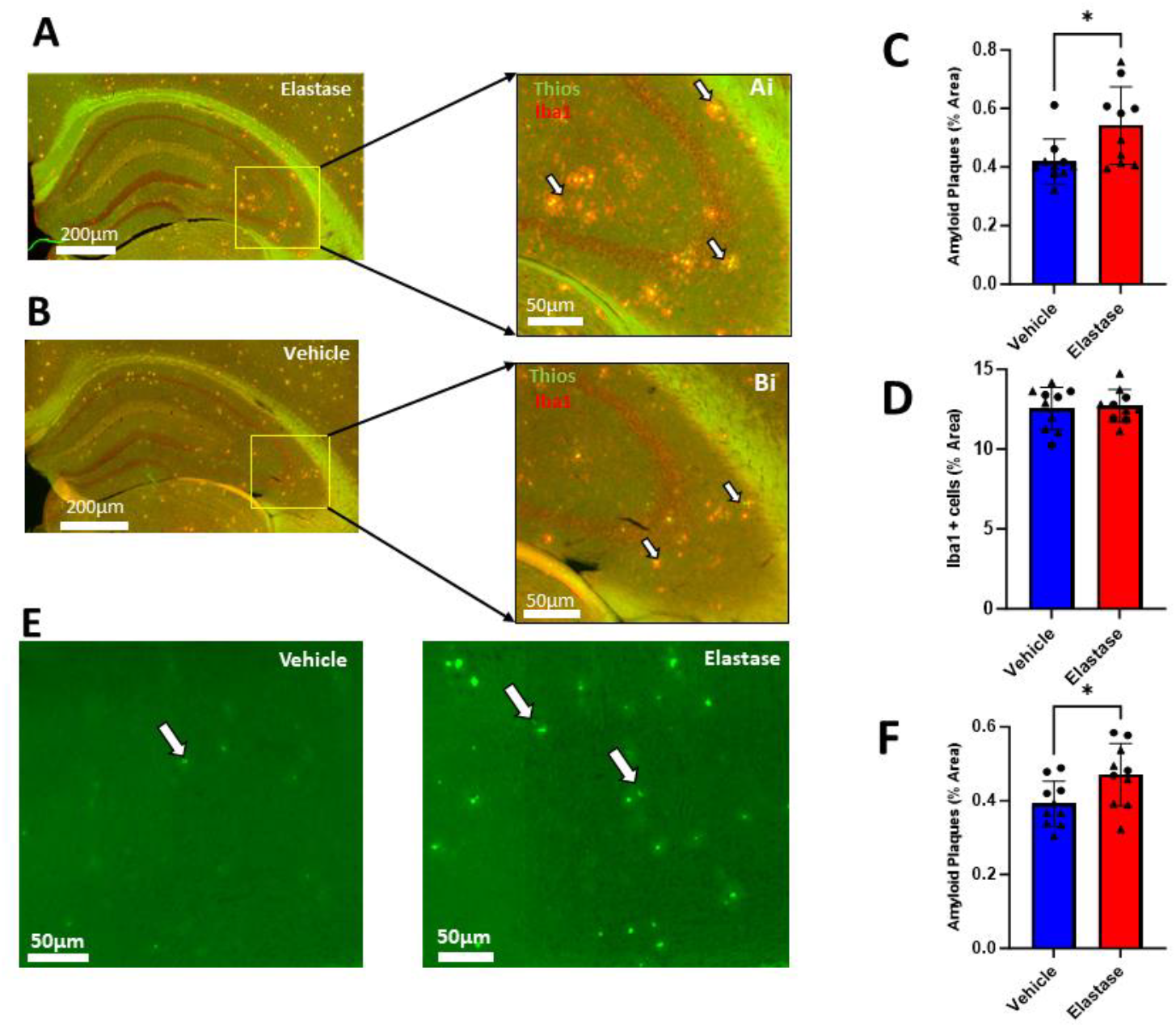
Dolichoectasia induces amyloid plaque deposition in the hippocampus without triggering neuroinflammation. **[A & B]** Amyloid plaque pathology and microglial activation were assessed in the hippocampus of App^NL-G-F^ mice by immunostaining with Thioflavin-S (ThioS) to label amyloid-β plaques and Iba1 to label activated microglial cells. ThioS+/Iba1+ staining in the hippocampus of an elastase-injected App^NL-G-F^ mouse. **[Ai & Bi]** Positive signals of amyloid plaque (green) and Iba1 (red) were seen throughout the hippocampus. The microglial cells can be seen engulfing the plaque (yellow) with an effort in trying to remove it from the hippocampus. **[C]** Elastase increased the accumulation of amyloid plaque in the hippocampus, [t(18)= 2.540, **P= 0.021 &** F(1,16)= 8.753, **P= 0.009 r**espectively; n= 10 per group]. **[C]** An overall sex difference was seen between groups, [F(1,16)= 6.370, **P= 0.023];** male: n=6 and female: n=4 per group. Female *App*^*NL-G-F*^ mice were trending to significance regarding the effect of elastase promoting the accumulation of amyloid plaque, [P> 0.058]; n=4 per group. **[D]** There was no increase in neuroinflammation (Iba1 activation) in the total hippocampus after elastase injection [**t**(18)= 0.303, p= 0.765; n=10 per group]; however, looking at the CA3 a trend towards increased microglial activation was seen in male mice, [P= 0.092]; n=6 per group. **[E]** In the cortex, elastase causes an increase in amyloid plaque deposition [F] t(18)= 2.378, **P= 0.029 &** F(1,16)= 7.358, **P= 0.015**, respectively; n=10 per group]. **[F]** An overall sex difference was also seen between groups, F(1,16)= 17.96, **P= 0.0006;** male: n=6 and female: n=4 per group. A significant difference in increase amyloid deposition was seen in the male *App*^*NL-G-F*^ mice with elastase in comparison to vehicle treated mice **[P= 0.023]**. An unpaired t-test was done to compare elastase and vehicle groups. Additionally, a two-way ANOVA with Sidak’s post hoc test for multiple comparisons was done to analyze the effects of sex. ● Female and ▲ Male

While we did not observe an increase in neuroinflammation throughout the hippocampus alongside elevated amyloid plaque deposition, a trend towards increased neuroinflammation was evident in the CA3 region of elastase-treated male mice compared to vehicle-treated male mice (p=0.093). We then evaluated the activation of astrocytes in the hippocampus of *App*^*NL-G-F*^ mice. Immunohistochemistry staining astrocytes using the GFAP antibody, did not demonstrate a difference in astrocyte activation in 9-month-old *App*^*NL-G-F*^ mice between treated and control groups [t (18) = 1.231, **p= 0.234**, F(1,16)= 1.968, p= 0.180]. We further tested if the CA1 or CA3 regions had early signs of astrogliosis and found no evidence of upregulated astrocytic activation in either region. These results also suggest that elastase may not play a role in sustaining astrocyte activation, or the response was attenuated over time as elastase administration was done 3 months prior. As evidence indicates that dolichoectasia resulted in increased plaque deposition, we do not exclude that a cascade of events is activated at the time of elastase induction, including neuroinflammation.

### Quantification of MMP-9 protein in the cortex of *App*^*NL-G-F*^ mice with Dolichoectasia

In light of increased amyloid plaque accumulation without concurrent neuroinflammation, we proceeded to investigate possible underlying mechanisms driving this event. The focus on MMP-9 stems from its known activation with dolichoectasia (Gutierrez et al., 2016; Lopez-Navarro, Park, Willey, & Gutierrez, 2022; Pico et al., 2010; Zhu, Xing, Dai, Kallmes, & Kadirvel, 2017), likely due to its role in elastin degradation within blood vessels (X. Wang & Khalil, 2018) and disruption of the BBB (Kim & Hwang, 2011; Rosenberg, 2009). This disruption can increase BBB permeability allowing toxins to enter the brain. We conducted Western blot analysis on the cortex of *App*^*NL-G-F*^ mice to assess MMP-9 levels, given its emerging significance as a potential contributor to Alzheimer’s disease pathophysiology. Our focus was primarily on the MMP-9 active form at 67 kDa, with the pro-form at 92 kDa serving as a reference (**Fig. 2A**). Our findings demonstrated a strong trend in upregulation of the active form of MMP-9 in elastase-injected mice relative to vehicle [[t (18) = 2.029, **P= 0.058 &** F (1, 15) = 4.490, **P= 0.05**; **Fig. 2B]**, suggesting a potential role in the pathophysiology of AD.

**Figure 2:**
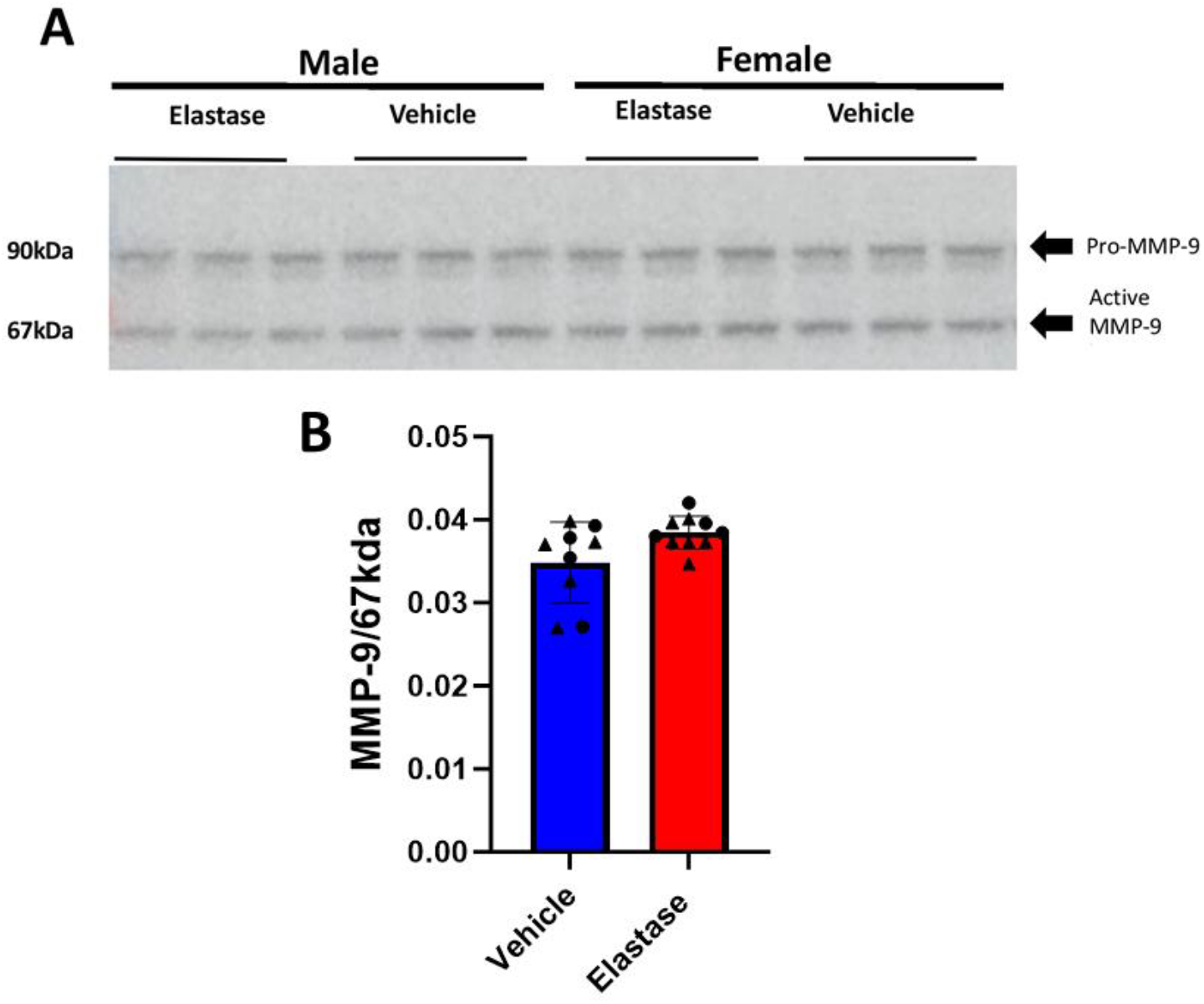
Quantification of MMP-9 protein in the cortex of 9 months old *App*^*NL-G-F*^ mice with Dolichoectasia. **[A]** Representative image of pro-mmp-9/92kda (upper band) and active mmp-9/67kda (lower band). **[B]** The quantification results of active MMP-9/67 kDa, normalized to the control GAPDH, show a trend towards increased MMP-9/67 kDa activity in the elastase-treated group compared to the vehicle group, [t (18) = 2.029, P= 0.058 & F (1, 15) = 4.490, P= 0.051; elastase (n=10) and vehicle (n=9), respectively]. An unpaired t-test was done to compare elastase and vehicle groups. Additionally, a two-way ANOVA with Sidak’s post hoc test for multiple comparisons was done to analyze the effects of sex. • Female and ▲ Male

### Elastase injection causes an increase in p62 expression showing a dysfunctional autophagy system

Given the observed increase in cortical MMP-9 levels in elastase-injected *App* ^NL-G-F^ mice, we next investigated whether autophagic function was altered, as matrix metalloproteinase activity has been linked to regulation of autophagy. Prior studies have shown that inhibition of MMPs upregulates autophagy-related genes (ATG5 and ATG7), both critical for autophagy initiation (Jo et al., 2011; Nandi et al., 2020). To assess autophagic-lysosomal flux in the context of AD-related pathology, we examined p62, a scaffold protein that binds ubiquitinated aggregates destined for degradation via the autophagic-lysosomal system (Li, Li, & Wu, 2022). An increase in p62 levels would indicate incomplete proteostasis. In our observations, we detected a significantly elevated level of p62 in the hippocampus following elastase injection [t (18) = 2.90, **p=0.009** and F(1,16) = 6.822, **p=0.019; Fig. 3B**] where elastase-injected male mice had a significantly higher p62 puncta burden than vehicle treated mice (**p=0.024**). In the cortex (**Fig. 3C**), the *App*^*NL-G-F*^ mice also have significantly higher levels of p62 protein following elastase injection [t (18) = 3.79, **P= 0.001 &** F(1,16)= 12.04, **P=0.003; Fig. 3D**]. Similarly, as in the hippocampus, the male elastase mice have greater p62 burden than vehicle male mice (**p=0.016**).

**Figure 3:**
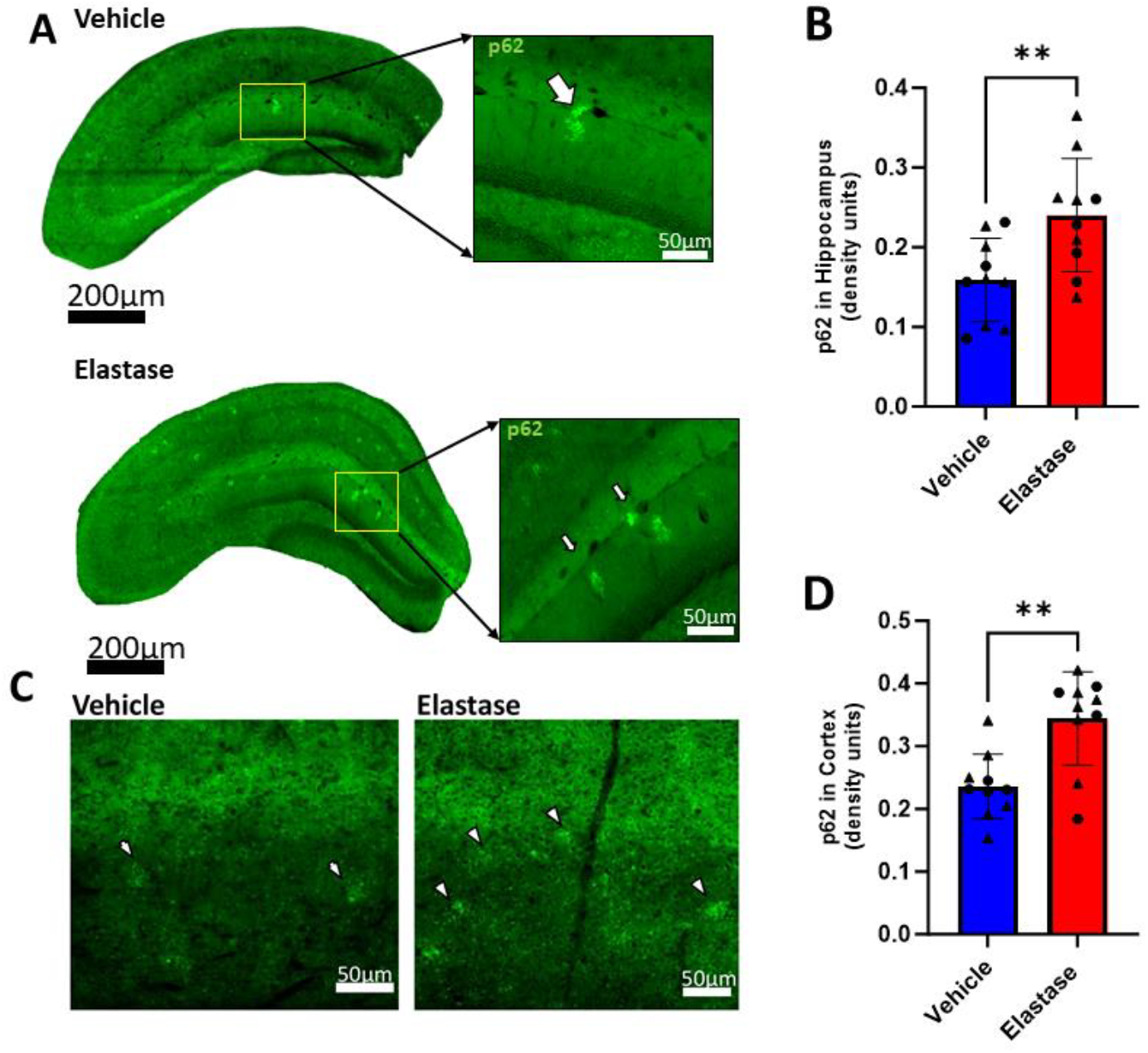
Elastase causes an increase in p62 expression highlighting the importance of the autophagy system in AD. To determine if autophagy is dysfunctional in *App*^*NL-G-F*^ mice we quantified the number of p62 clusters in the hippocampus. **[A]** p62+ staining in the hippocampus of elastase-injected male *App*^*NL-G-F*^ mice in comparison to PBS-injected male *App*^*NL-G-F*^ mice. The positive signal of p62 (green) can be seen as clusters (shown by a white arrow) in the hippocampus. The elastase *App*^*NL-G-F*^ mice were shown to have a greater amount of p62 clusters, compared to control mice. **[B]** Elastase increased the number of clusters in the hippocampus, [t (18) = 2.90, **P= 0.009 &** F (1, 16) = 6.822, **P= 0.019**; n= 10 per group]. Male *App*^*NL-G-F*^ mice are more susceptible to this increase in p62 clusters in the hippocampus as it pertains to the effect of elastase when compared vehicle treated male mice, [**P= 0.024**; n= 6 per group]. **[C]** Increased levels of p62 were also seen in the cortex after elastase injection [D] [t (18) = 3.79, **P= 0.001 &** F(1,16)= 12.04, **P= 0.003**; n= 10 per group] and male *App*^*NL-G-F*^ mice with elastase were shown to have more p62 clusters than vehicle treated male mice [**P= 0.016**; n= 6 per group]. An unpaired t-test was done to compare elastase and vehicle groups. Additionally, a two-way ANOVA with Sidak’s post hoc test for multiple comparisons was done to analyze the effects of sex. ● Female and ▲ Male

### Elastase-induced hippocampal neuron loss in *App*^*NL-G-F*^ mice

Another hallmark of AD is neuronal loss, and we assessed this alongside increased amyloid plaque deposition, in the hippocampus, an area central to learning and memory. Our observations reveal a significant neuronal loss in the cell layers of the hippocampus in App^NL-G-F^ mice following elastase injection. Density analysis shows no neuronal loss in the dentate gyrus (DG) (**Fig. 4A**). We then proceeded to specifically examine the CA1 and CA3 areas and observed neuronal loss, as measured by the mature neuron marker, NeuN, in elastase-injected mice, specifically in the CA3 region (**Fig. 4B**). The quantitative analysis further supports this observation, indicating that PBS-injected mice have more neurons than those injected with elastase, with a statistical trend toward significance [F(1,16)= 4.281, **P= 0.051; Fig. 4B**]. Female mice are more susceptible to the effects of elastase, exhibiting a more substantial neuronal loss compared to vehicle treated female mice (**p=0.017**). Similarly, in the CA1 region, a comparable loss of neuronal cells was evident (**Fig. 4C**), with an overall reduction in neuronal count in elastase-injected mice compared to vehicle (PBS control) [t (18) = 2.497, **p= 0.022 &** F(1,16) = 5.239, **p=0.0036; Fig 2C)**. A trend was also seen indicating that male mice might be more susceptible to neuronal loss in the CA1 region (p=0.085). Neuronal loss in the CA1 and CA3 regions is a classical feature in Alzheimer’s disease. (Padurariu, Ciobica, Mavroudis, Fotiou, & Baloyannis, 2012), vascular dementia and mixed dementia (Gemmell et al., 2012). In summary, our observations showed that elastase-injection in the App ^NL-G-F^ mice can lead to neuronal cell loss.

**Figure 4:**
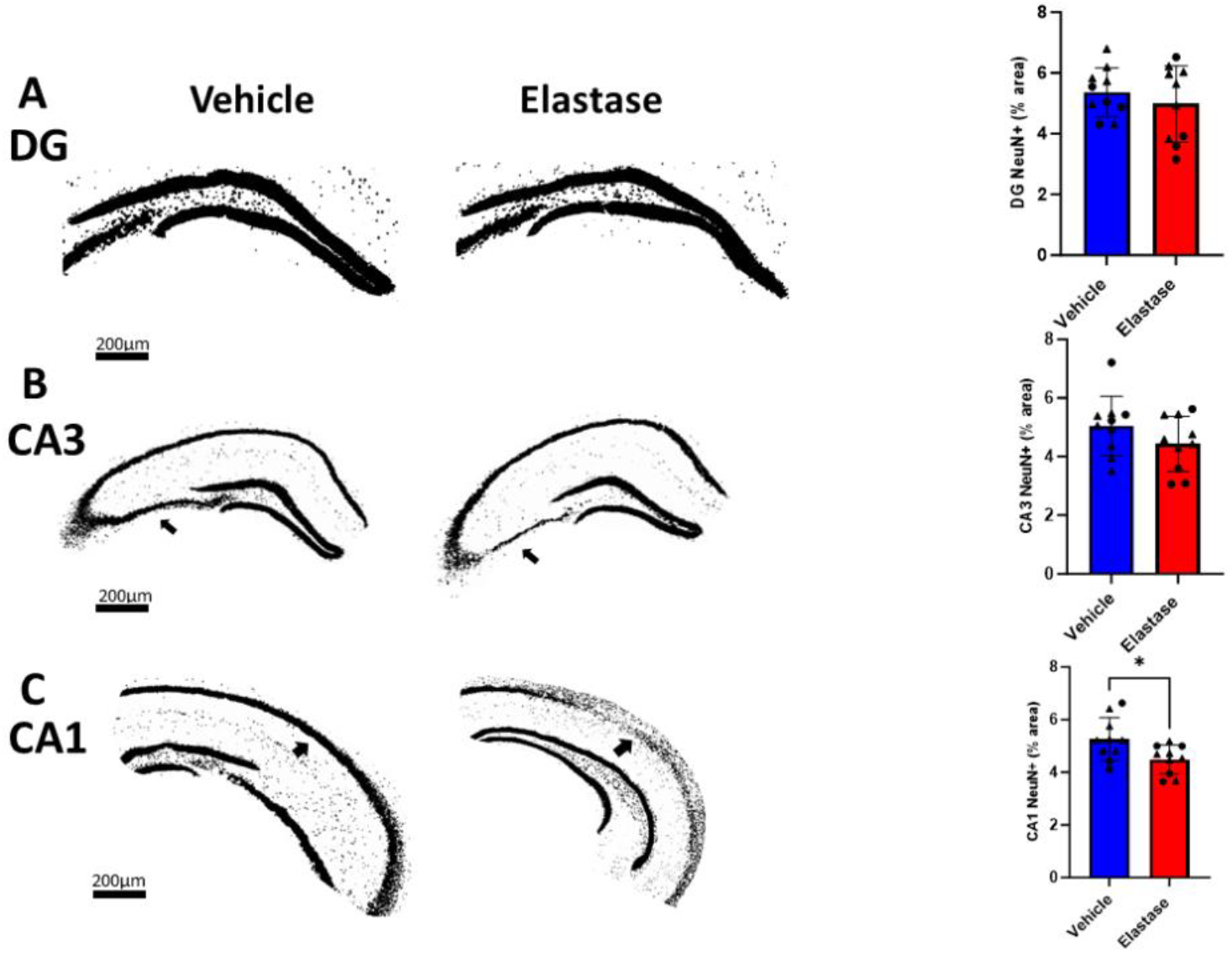
*App*^*NL-G-F*^ mice with dolichoectasia have neuronal loss in the hippocampus. To determine if elastase injection contributes to the loss of hippocampal neurons, we determined neuronal density in comparison to controls. **[A]** No neuronal loss was seen in the DG cell layer of *App*^*NL-G-F*^ mice after elastase injection [t (18) = 0.808, P= 0.429 & F (1, 16) = 0.884, P= 0.361; n=10 per group]. However, a trend towards significance for the overall sex difference was seen **[F (1, 16) = 4.111**, P > 0.060]. [**B**] The CA3 cell layer of *App*^*NL-G-F*^ mice displays a visual loss of NeuN (black) staining after elastase injection, compared to vehicle mice. Elastase might have a role in decreasing the density of NeuN staining in the cell layer of CA3 where an overall trend towards significance was seen, [t (18) = 1.411, P= 0.175 & F (1, 16) = 4.281, P> 0.055; n=10 per group]. The density of NeuN staining in the CA3 cell layer of female *App*^*NL-G-F*^ mice is significantly decreased after elastase injection, compared to vehicle [**P= 0.017**; n=4 per group]. **[C]** The CA1 cell layer of *App*^*NL-G-F*^ mice displays a significant loss of NeuN (black) staining after elastase injection, compared to vehicle mice. Elastase causes an overall decrease in the density of NeuN staining in the CA1 cell layer, [t (18) = 2.497, **P= 0.022 &** F (1, 16) = 5.239, **P= 0.036;** n= 10 per group]. Post-hoc analysis separating sex shows a trend towards neuronal loss in elastase male *App*^*NL-G-F*^ mice vs vehicle treated male mice [P = 0.085, n = 6 per group] and no change in female *App*^*NL-G-F*^ mice, suggesting that male mice might be more prone to the effects of elastase. An unpaired t-test was done to compare elastase and vehicle groups. Additionally, a two-way ANOVA with Sidak’s post hoc test for multiple comparisons was done to analyze the effects of sex. ● Female and ▲ Male

### Quantification of PHF1 protein in the cortex of *App*^*NL-G-F*^ mice with Dolichoectasia

In addition to plaques, neuroinflammation and neuronal loss, tauopathy represents another major AD-related phenotype. While these mice express human APP but not human tau, we measured hyperphosphorylated tau targeting the PHF1 (S396/T404 epitope) in the cortex of *App*^*NL-G-F*^ mice via Western blot analysis using the PHF1 antibody. Our analysis did not reveal any significant increase in hyperphosphorylated tau levels in elastase-injected mice compared to vehicle [t (18) = 0.007, P= 0.994 & F(1,16)= 0.0003, P= 0.986; **Fig. 5B**]. However, a notable overall sex effect mice [F (1, 16) = 9.051, P= **0.008; Fig. 5B]** was observed where both elastase and vehicle male mice exhibited higher levels of hyperphosphorylated tau than female.

**Figure 5:**
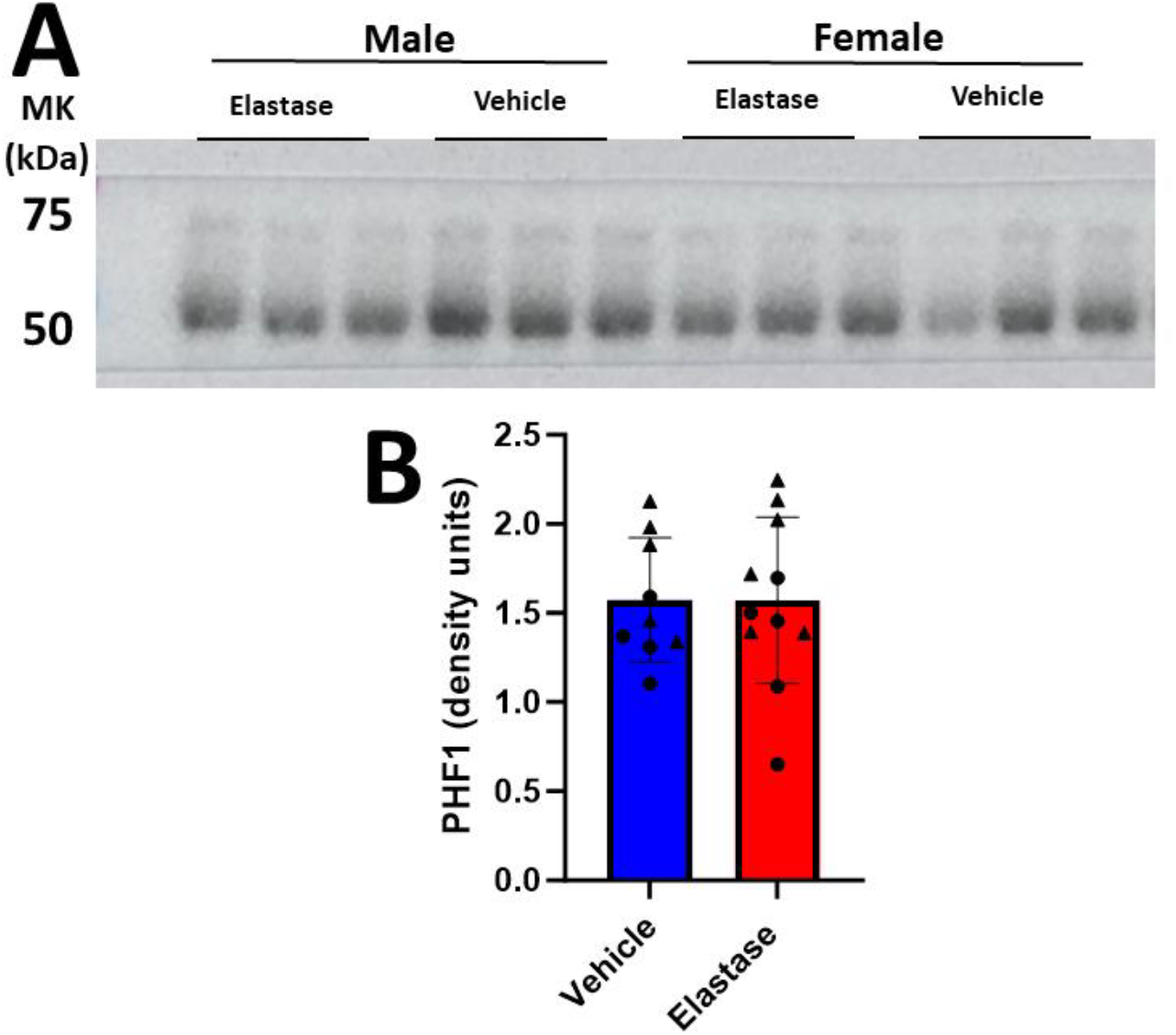
Quantification of PHF1 protein in the cortex of *App*^*NL-G-F*^ mice. **[A]** A representative image showing PHF1 bands across groups. **[B]** The quantification results of PHF1, normalized to the control GAPDH, where no significant difference was seen between groups [t (18) = 0.007, P= 0.994 & F (1, 16) = 0.0003, P= 0.986; elastase (n=11) and vehicle (n=9)]. However, a significant sex effect was seen, [F (1, 16) = 9.051, **P= 0.008;** male, elastase (n=6), vehicle (n=5); female, elastase (n=5), vehicle (n=4)]. The difference was between the male elastase mice having more PHF1 than the female elastase mice **[P= 0.043**, elastase (n=6); female, elastase (n=5)]. An unpaired t-test was done to compare elastase and vehicle groups. Additionally, a two-way ANOVA with Sidak’s post hoc test for multiple comparisons was done to analyze the effects of sex. ● Female and ▲ Male

## Discussion

The relevance of dolichoectasia/brain large artery dilatation and AD is an under-recognized and under-researched discipline, despite relating to 10-15% of dementia cases (Melgarejo et al., 2024). For this reason, we investigated the impact of experimental dolichoectasia induced by elastase to determine if it accelerates relevant pathological phenotypes seen in AD in an amyloid mouse model (App^NL-G-F^) (Dai et al., 2015; Simpson et al., 2024). The *App*^*NL-G-F*^ model from this study, mirrors amyloid plaque expression at physiological levels, aligning closely with early age-related Alzheimer’s disease phenotypes (Saito et al., 2014). This makes the AppNL-G-F model ideal for longitudinal studies, contrasting with other models characterized by amyloid plaque overexpression (Hsiao et al., 1996; Oakley et al., 2006). Our goal was to explore the effect of dolichoectasia on promoting AD pathology and to accomplish this, we opted to employ 6-month-old App^NL-G-F^, which has early amyloid pathology, as our model, administering elastase into the cisterna magna, located directly above the circle of Willis, which is composed of the most proximal and largest brain arteries (Simpson et al., 2024). Our study examined the impact of brain large artery dilatation by injecting elastase into the cisterna magna of *App*^*NL-G-F*^ mice at 6 months of age and allowing them to age to 9 months to model early AD-like changes.

Our findings are the first report to demonstrate than in amyloid plaque accumulation occurs with large artery dilatation, providing a causal relationship between dolichoectasia and AD pathology. A major goal of this work was to establish if there are potential pathological drivers related to arterial dilatation. In this study, we identified MMP-9 as a potential contributor as evidenced by increased MMP-9 in brains of elastase treated mice.

While increased amyloid plaque typically triggers neuroinflammation through microglial activation (C. Wang et al., 2023), our study did not observe an overall increase. We hypothesize that this discrepancy might arise from the 3-month interval after elastase injection. Elastase was administered to 6-month-old mice, which were then allowed to age until 9 months before being sacrificed. It is possible that inflammation induced by elastase could have appeared earlier and subsequently resolved by the 9-month time point. Subsequent analysis of specific hippocampal regions (CA3) revealed a trend toward neuroinflammation in male elastase mice, though its persistence or progressive nature remains debatable. In addition, we observed no increase in astrogliosis, despite reported upregulation in astrocytic activation in tandem with microglial activation (Bouvier et al., 2016; Perez-Nievas & Serrano-Pozo, 2018). However, we did visually see increased IBA-1 immunoreactivity surrounding plaques, suggesting a more nuanced increase, which requires more mechanistic exploration.

We observed an overall increase in MMP-9 expression in the cortex of elastase-injected mice compared to vehicle. This upregulation correlates with previous findings linking elastase to elevated MMP expression (Ferry et al., 1997; Garratt et al., 2015). MMPs are widely expressed by astrocytes (Muir et al., 2002), macrophages (Newby, 2008), and microglial cells (Lassmann, Zimprich, Vass, & Hickey, 1991) and play pivotal roles in extracellular matrix breakdown and remodeling. In this study, our primary focus was on MMP-9 due to its role in the pathophysiology of Alzheimer’s disease (Asahina, Yoshiyama, & Hattori, 2001; Lorenzl et al., 2003). Elevated expression of MMP-9 is associated with potential disruptions in the blood-brain barrier (Sandoval & Witt, 2008; Svedin, Hagberg, Savman, Zhu, & Mallard, 2007), dilatation of blood vessels (aka dolichoectasia) (Gutierrez et al., 2016; Zhu et al., 2017), and the promotion of neuroinflammation (Dwir et al., 2020).

The increased MMP-9 expression prompted further investigation into its potential pathological roles in AD. Initially, we hypothesized that elastase-injected mice might exhibit dysfunctional autophagy, as prior research suggests MMP-9 and MMP-2 inhibition leads to increased autophagy (Jo et al., 2011; Nandi et al., 2020). This prompted us to explore whether there might be an MMP-9 upregulation that correlates with autophagic dysfunction. Our findings in the hippocampus and cortex of *App*^*NL-G-F*^ elastase mice revealed elevated p62 expression, typically forming clusters, indicative of impaired autophagic-lysosomal system function (Jiang & Mizushima, 2015). This disruption prevents the removal of abnormal proteins, such as amyloid plaque, from the brain by impeding their degradation (Lee et al., 2022). Consequently, this disruption exposes the brain to the harmful effects of accumulated abnormal proteins, elevating the risk of neuronal loss (J. Liu & Li, 2019). Confirming this hypothesis, we observed increased amyloid plaque in the hippocampus and cortex of elastase mice through Thioflavin S staining, further reinforcing the connection between dysfunctional autophagy and AD pathology.

The prosepct aht f MMP-9 disrupts autophagy was further supported by previous studies showing inhibiting MMPs causes an upregulation of the genes ATG5 (MMP-2) and ATG7 (MMP-9), which are crucial for the autophagic process (Jo et al., 2011; Nandi et al., 2020).). The accumulation of p62 that was observed in our studies suggesting protein clearance may be defective with arterial dilatation. Further, this confirms past findings that when autophagy is altered by increased autophagosomes, it has a potential to increase amyloid production (Nixon et al., 2005; Yu et al., 2005). Further, a loss of mature neurons in AD (Niikura, Tajima, & Kita, 2006) may be related to faulty protein clearance and increased amyloid burden, which we observed in the CA1 and CA3 layers of the hippocampus, an area severely impacted in Alzheimer’s, and associated the learning and memory. This aligns with established literature documenting neuronal loss in CA1 and CA3 regions in AD (Padurariu et al., 2012), suggesting that neuroinflammation following elastase injection might inflict permanent neuronal damage, then subsequently resolving or dampening. Additionally, amyloid plaque and tau hyperphosphorylation accumulation, may also contribute to neuronal degeneration (Rajmohan & Reddy, 2017).

Finally, we saw no increase in the hyperphosphorylation of tau after elastase injection using the PHF1 antibody that recognizes tau protein phosphorylation at Ser396 and Ser404 residues (Otvos et al., 1994). Despite these mice primarily modeling amyloid plaque, we predicted that there might be some degree of tau hyperphosphorylation. Cortical analysis revealed no difference in PHF1 expression between elastase and vehicle groups, but a notable sex difference emerged, with male mice exhibiting higher expression than female mice, possibly attributed to the neuroprotective effects of estrogen or progesterone in females (Carroll et al., 2007).

In addition to upregulating MMP-9 expression, elastase may exacerbate AD pathology through structural brain changes. Elastase-induced blood vessel dilation and elongation (Dai et al., 2015), diminishes cerebral blood flow and can be detrimental to neurons and the blood-brain barrier (de la Torre & Mussivand, 1993). BBB disruption exposes the central nervous system (CNS) to peripheral immune cells from the bloodstream, which could cause an increase in neuroinflammation. Furthermore, this breach permits the entry of amyloid beta (Aβ) peptides into the brain, which contribute to amyloid plaque accumulation (Acharya et al., 2024). Another mechanism for BBB disruption involves the relocation of tight junction proteins away from the BBB to repair damaged blood vessels, thereby weakening its integrity (Biron et al., 2011).

However, BBB disruption represents just one avenue for substances to infiltrate the brain. In response to defective blood vessels, the body initiates compensatory angiogenesis, mediated by VEGF. While this process typically aims to generate new blood vessels, in cases of excessive VEGF expression, pathological angiogenesis may occur. This results in the formation of immature, leaky blood vessels, facilitating the entry of fluids into the brain interstitial space (Desai et al., 2009).

In summary, our study reveals that dolichoectasia, characterized by the dilatation of large blood vessels, can exacerbate AD pathology. We demonstrated that dolichoectasia induces an upregulation of MMP-9 expression in the cortex of App^NL-G-F^ mice. This upregulation in MMP-9 is believed to disrupt autophagy, evidenced by elevated p62 levels in both the hippocampus and cortex, particularly pronounced in male mice. We hypothesize that this disruption in autophagy leads to increased amyloid plaque accumulation in both regions. Males exhibit higher cortical plaque levels and p62 levels in the hippocampus and cortex, suggesting that males are more susceptible to the effects of dolichoectasia. Furthermore, elevated amyloid plaque levels may contribute to amyloid toxicity, resulting in neuronal loss in the CA1 and CA3 cell layers, with males and females experiencing neuronal loss. These findings show the differential impact of dolichoectasia on male and female mice, highlighting its influence on AD pathology. Potential future studies include behavioral testing for cognitive impairment and earlier immunological staining for microglial activation post-elastase injection.

## ABBREVIATIONS

AD: alzheimer’s disease
Aβ: amyloid beta
APP: amyloid precursor protein
BBB: blood-brain barrier
CNS: central nervous system
CSF: cerebrospinal fluid
CM: cisterna magna
DG: dentate gyrus
DTT: dithiothretiol
MMP: matrix metalloproteinases
NSAID: nonsteroidal anti-inflammatory drug
TBS-T: tris buffered saline 0.05% tween 20
VaD: vascular dementia
VEGF: vascular endothelial growth factor

## Author’s contribution

**Dominic Simpson**: Writing – original draft; review & editing; formal analysis; investigation; visualization. **Wai Haung Yu**: Writing – original draft; review & editing; conceptualization; funding; methodology; project administration; resources; supervision. **Christopher D. Morrone**: Writing –review & editing; methodology. **Darcy Wear**: Writing –review & editing. **Adrian Khani**: Investigation. **Fang Liu**: Writing – review & editing. **Jose Guttierez**: Writing – review & editing; funding, conceptualization.

## Acknowledgments

This research was supported by the National Institutes of Health (AG066162).

## Disclosure

There is no conflict of interest.

---

**Supplemental Table 1:** Primary antibodies used in this study.

| **Primary Antibodies** | **Vendor** | **Catalog #** | **Dilution** |
| --- | --- | --- | --- |
| GFAP | Dako | Z0334 | 1:500 |
| Iba1 | Wako | 019-19741 | 1:1000 |
| P62 | Abcam | Ab109012 | 1:400 |
| NeuN | Millipore Sigma | MAB377 | 1:500 |
| Thioflavin S | Millipore Sigma | T1892 | - |
| MMP-9 | Proteintech | 10375-2-AP | 1:1500 |
| Phf1 | Gift from Peter Davies | - | 1:1000 |
| GAPDH | Sigma-Aldrich | G8795 | 1:4000 |

| **Secondary Antibodies** | **Vendor** | **Catalog #** | **Dilution** |
| --- | --- | --- | --- |
| **Immunohistochemistry** |  |  |  |
| Goat anti-rabbit 488 | Invitrogen | A11008 | 1:200 |
| Goat anti-rabbit 568 | Invitrogen | A11075 | 1:200 |
| Goat anti-mouse 568 | Invitrogen | A11004 | 1:200 |
| **Biochemistry** |  |  |  |
| horseradish peroxidase-Conjugated goat anti-mouse IgG | Jackson ImmunoResearch<br>Labs | 115-035-003 | 1:10000 |
| horseradish peroxidase-Conjugated goat anti-rabbit IgG | Jackson ImmunoResearch<br>Labs | 111-035-008 | 1:10000 |

## Notes

### Competing Interest Statement

The authors have declared no competing interest.

## References

Acharya, N. K., Grossman, H. C., Clifford, P. M., Levin, E. C., Light, K. R., Choi, H., … Nagele, R. G. (2024). A Chronic Increase in Blood-Brain Barrier Permeability Facilitates Intraneuronal Deposition of Exogenous Bloodborne Amyloid-Beta1-42 Peptide in the Brain and Leads to Alzheimer’s Disease-Relevant Cognitive Changes in a Mouse Model. J Alzheimers Dis, 98(1), 163–186. doi:10.3233/JAD-231028

Asahina, M., Yoshiyama, Y., & Hattori, T. (2001). Expression of matrix metalloproteinase-9 and urinary-type plasminogen activator in Alzheimer’s disease brain. Clin Neuropathol, 20(2), 60–63. Retrieved from https://www.ncbi.nlm.nih.gov/pubmed/11327298

Barker, W. W., Luis, C. A., Kashuba, A., Luis, M., Harwood, D. G., Loewenstein, D., … Duara, R. (2002). Relative frequencies of Alzheimer disease, Lewy body, vascular and frontotemporal dementia, and hippocampal sclerosis in the State of Florida Brain Bank. Alzheimer Dis Assoc Disord, 16(4), 203–212. doi:10.1097/00002093-200210000-00001

Biron, K. E., Dickstein, D. L., Gopaul, R., & Jefferies, W. A. (2011). Amyloid triggers extensive cerebral angiogenesis causing blood brain barrier permeability and hypervascularity in Alzheimer’s disease. PLoS One, 6(8), e23789. doi:10.1371/journal.pone.0023789

Bouvier, D. S., Jones, E. V., Quesseveur, G., Davoli, M. A., T, A. F., Quirion, R., … Murai, K. K. (2016). High Resolution Dissection of Reactive Glial Nets in Alzheimer’s Disease. Sci Rep, 6, 24544. doi:10.1038/srep24544

Brandl, S., & Reindl, M. (2023). Blood-Brain Barrier Breakdown in Neuroinflammation: Current In Vitro Models. Int J Mol Sci, 24(16). doi:10.3390/ijms241612699

Carroll, J. C., Rosario, E. R., Chang, L., Stanczyk, F. Z., Oddo, S., LaFerla, F. M., & Pike, C. J. (2007). Progesterone and estrogen regulate Alzheimer-like neuropathology in female 3xTg-AD mice. J Neurosci, 27(48), 13357–13365. doi:10.1523/JNEUROSCI.2718-07.2007

Cortes-Canteli, M., & Iadecola, C. (2020). Alzheimer’s Disease and Vascular Aging: JACC Focus Seminar. J Am Coll Cardiol, 75(8), 942–951. doi:10.1016/j.jacc.2019.10.062

Dai, D., Kadirvel, R., Rezek, I., Ding, Y. H., Lingineni, R., & Kallmes, D. (2015). Elastase-induced intracranial dolichoectasia model in mice. Neurosurgery, 76(3), 337–343; discussion 343. doi:10.1227/NEU.0000000000000615

de la Torre, J.C., & Mussivand, T. (1993). Can disturbed brain microcirculation cause Alzheimer’s disease? Neurol Res, 15(3), 146–153. doi:10.1080/01616412.1993.11740127

Del Brutto, V. J., Ortiz, J. G., & Biller, J. (2017). Intracranial Arterial Dolichoectasia. Front Neurol, 8, 344. doi:10.3389/fneur.2017.00344

Desai, B. S., Schneider, J. A., Li, J. L., Carvey, P. M., & Hendey, B. (2009). Evidence of angiogenic vessels in Alzheimer’s disease. J Neural Transm (Vienna), 116(5), 587–597. doi:10.1007/s00702-009-0226-9

DeTure, M. A., & Dickson, D. W. (2019). The neuropathological diagnosis of Alzheimer’s disease. Mol Neurodegener, 14(1), 32. doi:10.1186/s13024-019-0333-5

Duong, S., Patel, T., & Chang, F. (2017). Dementia: What pharmacists need to know. Can Pharm J (Ott), 150(2), 118–129. doi:10.1177/1715163517690745

Dwir, D., Giangreco, B., Xin, L., Tenenbaum, L., Cabungcal, J. H., Steullet, P., … Do, K. Q. (2020). MMP9/RAGE pathway overactivation mediates redox dysregulation and neuroinflammation, leading to inhibitory/excitatory imbalance: a reverse translation study in schizophrenia patients. Mol Psychiatry, 25(11), 2889–2904. doi:10.1038/s41380-019-0393-5

Ferry, G., Lonchampt, M., Pennel, L., de Nanteuil, G., Canet, E., & Tucker, G. C. (1997). Activation of MMP-9 by neutrophil elastase in an in vivo model of acute lung injury. FEBS Lett, 402(2-3), 111–115. doi:10.1016/s0014-5793(96)01508-6

Garratt, L. W., Sutanto, E. N., Ling, K. M., Looi, K., Iosifidis, T., Martinovich, K. M., … Australian Respiratory Early Surveillance Team for Cystic, F. (2015). Matrix metalloproteinase activation by free neutrophil elastase contributes to bronchiectasis progression in early cystic fibrosis. Eur Respir J, 46(2), 384–394. doi:10.1183/09031936.00212114

Gemmell, E., Bosomworth, H., Allan, L., Hall, R., Khundakar, A., Oakley, A. E., … Kalaria, R. N. (2012). Hippocampal neuronal atrophy and cognitive function in delayed poststroke and aging-related dementias. Stroke, 43(3), 808–814. doi:10.1161/STROKEAHA.111.636498

Gutierrez, J., Guzman, V., Khasiyev, F., Manly, J., Schupf, N., Andrews, H., … Brickman, A. M. (2019). Brain arterial dilatation and the risk of Alzheimer’s disease. Alzheimers Dement, 15(5), 666–674. doi:10.1016/j.jalz.2018.12.018

Gutierrez, J., Menshawy, K., Goldman, J., Dwork, A. J., Elkind, M. S., Marshall, R. S., & Morgello, S. (2016). Metalloproteinases and Brain Arterial Remodeling Among Individuals With and Those Without HIV Infection. J Infect Dis, 214(9), 1329–1335. doi:10.1093/infdis/jiw385

Gutierrez, J., Sacco, R. L., & Wright, C. B. (2011). Dolichoectasia-an evolving arterial disease. Nat Rev Neurol, 7(1), 41–50. doi:10.1038/nrneurol.2010.181

Hsiao, K., Chapman, P., Nilsen, S., Eckman, C., Harigaya, Y., Younkin, S., … Cole, G. (1996). Correlative memory deficits, Abeta elevation, and amyloid plaques in transgenic mice. Science, 274(5284), 99–102. doi:10.1126/science.274.5284.99

Jiang, P., & Mizushima, N. (2015). LC3-and p62-based biochemical methods for the analysis of autophagy progression in mammalian cells. Methods, 75, 13–18. doi:10.1016/j.ymeth.2014.11.021

Jo, Y. K., Park, S. J., Shin, J. H., Kim, Y., Hwang, J. J., Cho, D. H., & Kim, J. C. (2011). ARP101, a selective MMP-2 inhibitor, induces autophagy-associated cell death in cancer cells. Biochem Biophys Res Commun, 404(4), 1039–1043. doi:10.1016/j.bbrc.2010.12.106

Kim, E. M., & Hwang, O. (2011). Role of matrix metalloproteinase-3 in neurodegeneration. J Neurochem, 116(1), 22–32. doi:10.1111/j.1471-4159.2010.07082.x

Lassmann, H., Zimprich, F., Vass, K., & Hickey, W. F. (1991). Microglial cells are a component of the perivascular glia limitans. J Neurosci Res, 28(2), 236–243. doi:10.1002/jnr.490280211

Lee, J. H., Yang, D. S., Goulbourne, C. N., Im, E., Stavrides, P., Pensalfini, A., … Nixon, R. A. (2022). Faulty autolysosome acidification in Alzheimer’s disease mouse models induces autophagic build-up of Abeta in neurons, yielding senile plaques. Nat Neurosci, 25(6), 688–701. doi:10.1038/s41593-022-01084-8

Li, Y., Li, S., & Wu, H. (2022). Ubiquitination-Proteasome System (UPS) and Autophagy Two Main Protein Degradation Machineries in Response to Cell Stress. Cells, 11(5). doi:10.3390/cells11050851

Liu, F. X., Niu, Y. G., Zhang, D. P., Zhang, H. L., Zhang, Z. Q., Sun, R. Q., & Zhang, Y. K. (2022). Modified Protocol for Establishment of Intracranial Arterial Dolichoectasia Model by Injection of Elastase Into Cerebellomedullary Cistern in Mice. Front Neurol, 13, 860541. doi:10.3389/fneur.2022.860541

Liu, J., & Li, L. (2019). Targeting Autophagy for the Treatment of Alzheimer’s Disease: Challenges and Opportunities. Front Mol Neurosci, 12, 203. doi:10.3389/fnmol.2019.00203

Livingston, G., Huntley, J., Liu, K. Y., Costafreda, S. G., Selbaek, G., Alladi, S., … Mukadam, N. (2024). Dementia prevention, intervention, and care: 2024 report of the Lancet standing Commission. Lancet, 404(10452), 572–628. doi:10.1016/S0140-6736(24)01296-0

Lopez-Navarro, E. R., Park, S., Willey, J. Z., & Gutierrez, J. (2022). Dolichoectasia: a brain arterial disease with an elusive treatment. Neurol Sci, 43(8), 4901–4908. doi:10.1007/s10072-022-06078-9

Lorenzl, S., Albers, D. S., Relkin, N., Ngyuen, T., Hilgenberg, S. L., Chirichigno, J., … Beal, M. F. (2003). Increased plasma levels of matrix metalloproteinase-9 in patients with Alzheimer’s disease. Neurochem Int, 43(3), 191–196. doi:10.1016/s0197-0186(03)00004-4

Melgarejo, J. D., Gurel, K., Compton, C. R., Liu, M., Guzman, V., Assuras, S., … Gutierrez, J. (2024). Brain artery diameters and risk of dementia and stroke. Alzheimers Dement, 20(4), 2497–2507. doi:10.1002/alz.13712

Muir, E. M., Adcock, K. H., Morgenstern, D. A., Clayton, R., von Stillfried, N., Rhodes, K., … Rogers, J. H. (2002). Matrix metalloproteases and their inhibitors are produced by overlapping populations of activated astrocytes. Brain Res Mol Brain Res, 100(1-2), 103-117. doi:10.1016/s0169-328x(02)00132-8

Nandi, S. S., Katsurada, K., Sharma, N. M., Anderson, D. R., Mahata, S. K., & Patel, K. P. (2020). MMP9 inhibition increases autophagic flux in chronic heart failure. Am J Physiol Heart Circ Physiol, 319(6), H1414–H1437. doi:10.1152/ajpheart.00032.2020

Newby, A. C. (2008). Metalloproteinase expression in monocytes and macrophages and its relationship to atherosclerotic plaque instability. Arterioscler Thromb Vasc Biol, 28(12), 2108–2114. doi:10.1161/ATVBAHA.108.173898

Niikura, T., Tajima, H., & Kita, Y. (2006). Neuronal cell death in Alzheimer’s disease and a neuroprotective factor, humanin. Curr Neuropharmacol, 4(2), 139–147. doi:10.2174/157015906776359577

Nixon, R. A., Wegiel, J., Kumar, A., Yu, W. H., Peterhoff, C., Cataldo, A., & Cuervo, A. M. (2005). Extensive involvement of autophagy in Alzheimer disease: an immuno-electron microscopy study. J Neuropathol Exp Neurol, 64(2), 113–122. doi:10.1093/jnen/64.2.113

Oakley, H., Cole, S. L., Logan, S., Maus, E., Shao, P., Craft, J., … Vassar, R. (2006). Intraneuronal beta-amyloid aggregates, neurodegeneration, and neuron loss in transgenic mice with five familial Alzheimer’s disease mutations: potential factors in amyloid plaque formation. J Neurosci, 26(40), 10129–10140. doi:10.1523/JNEUROSCI.1202-06.2006

Otvos, L., Jr., Feiner, L., Lang, E., Szendrei, G. I., Goedert, M., & Lee, V. M. (1994). Monoclonal antibody PHF-1 recognizes tau protein phosphorylated at serine residues 396 and 404. J Neurosci Res, 39(6), 669–673. doi:10.1002/jnr.490390607

Padurariu, M., Ciobica, A., Mavroudis, I., Fotiou, D., & Baloyannis, S. (2012). Hippocampal neuronal loss in the CA1 and CA3 areas of Alzheimer’s disease patients. Psychiatr Danub, 24(2), 152–158.Retrieved from https://www.ncbi.nlm.nih.gov/pubmed/22706413

Perez-Nievas, B. G., & Serrano-Pozo, A. (2018). Deciphering the Astrocyte Reaction in Alzheimer’s Disease. Front Aging Neurosci, 10, 114. doi:10.3389/fnagi.2018.00114

Pico, F., Jacob, M. P., Labreuche, J., Soufir, N., Touboul, P. J., Benessiano, J., … Amarenco, P. (2010). Matrix metalloproteinase-3 and intracranial arterial dolichoectasia. Ann Neurol, 67(4), 508–515. doi:10.1002/ana.21922

Rajmohan, R., & Reddy, P. H. (2017). Amyloid-Beta and Phosphorylated Tau Accumulations Cause Abnormalities at Synapses of Alzheimer’s disease Neurons. J Alzheimers Dis, 57(4), 975–999. doi:10.3233/JAD-160612

Rosenberg, G. A. (2009). Matrix metalloproteinases and their multiple roles in neurodegenerative diseases. Lancet Neurol, 8(2), 205–216. doi:10.1016/S1474-4422(09)70016-X

Saito, T., Matsuba, Y., Mihira, N., Takano, J., Nilsson, P., Itohara, S., … Saido, T. C. (2014). Single App knock-in mouse models of Alzheimer’s disease. Nat Neurosci, 17(5), 661–663. doi:10.1038/nn.3697

Sandoval, K. E., & Witt, K. A. (2008). Blood-brain barrier tight junction permeability and ischemic stroke. Neurobiol Dis, 32(2), 200–219. doi:10.1016/j.nbd.2008.08.005

Simpson, D., Morrone, C. D., Wear, D., Gutierrez, J., & Yu, W. H. (2024). A Visual Approach for Inducing Dolichoectasia in Mice to Model Large Vessel-Mediated Cerebrovascular Dysfunction. J Vis Exp(207). doi:10.3791/66792

Svedin, P., Hagberg, H., Savman, K., Zhu, C., & Mallard, C. (2007). Matrix metalloproteinase-9 gene knock-out protects the immature brain after cerebral hypoxia-ischemia. J Neurosci, 27(7), 1511–1518. doi:10.1523/JNEUROSCI.4391-06.2007

Wang, C., Zong, S., Cui, X., Wang, X., Wu, S., Wang, L., … Lu, Z. (2023). The effects of microglia-associated neuroinflammation on Alzheimer’s disease. Front Immunol, 14, 1117172. doi:10.3389/fimmu.2023.1117172

Wang, X., & Khalil, R. A. (2018). Matrix Metalloproteinases, Vascular Remodeling, and Vascular Disease. Adv Pharmacol, 81, 241–330. doi:10.1016/bs.apha.2017.08.002

Yu, W. H., Cuervo, A. M., Kumar, A., Peterhoff, C. M., Schmidt, S. D., Lee, J. H., … Nixon, R. A. (2005). Macroautophagy--a novel Beta-amyloid peptide-generating pathway activated in Alzheimer’s disease. J Cell Biol, 171(1), 87–98. doi:10.1083/jcb.200505082

Zenaro, E., Piacentino, G., & Constantin, G. (2017). The blood-brain barrier in Alzheimer’s disease. Neurobiol Dis, 107, 41–56. doi:10.1016/j.nbd.2016.07.007

Zhu, Y. Q., Xing, H., Dai, D., Kallmes, D. F., & Kadirvel, R. (2017). Differential Interstrain Susceptibility to Vertebrobasilar Dolichoectasia in a Mouse Model. AJNR Am J Neuroradiol, 38(3), 611–616. doi:10.3174/ajnr.A5028

